# Comprehensive epigenomic profiling of human alveolar epithelial differentiation identifies key epigenetic states and transcription factor co-regulatory networks for maintenance of distal lung identity

**DOI:** 10.1101/2021.04.14.439786

**Authors:** Beiyun Zhou, Theresa Ryan Stueve, Evan A. Mihalakakos, Lin Miao, Daniel J Mullen, Yinchong Wang, Yixin Liu, Jiao Luo, Evelyn Tran, Kim D. Siegmund, Sean K. Lynch, Amy L. Ryan, Ite A. Offringa, Zea Borok, Crystal N. Marconett

## Abstract

Disruption of alveolar epithelial cell (AEC) differentiation is implicated in peripheral lung diseases strongly impacting morbidity and mortality worldwide, such as chronic obstructive pulmonary disease, idiopathic pulmonary fibrosis, and lung adenocarcinoma. Elucidating underlying disease pathogenesis requires a mechanistic molecular understanding of AEC differentiation. However, to date no study has comprehensively characterized the dynamic epigenomic alterations that facilitate this critical process in humans. We comprehensively profiled the epigenomic states of human AECs during type 2 to type 1-like cell differentiation, including the methylome and chromatin functional domains, and integrated this with transcriptome-wide RNA expression. Enhancer regions were drastically altered during AEC differentiation. Transcription factor binding analysis within enhancer regions revealed diverse interactive networks with enrichment for dozens of transcription factors, including NKX2-1 and FOXA family members, as well as transcription factors with previously uncharacterized roles in lung differentiation, such as members of the MEF2, TEAD, and AP1 families. Additionally, associations between transcription factors changed during differentiation, implicating a complex network of heterotrimeric complex switching may be involved in facilitating differentiation. Integration of AEC enhancer states with the catalog of enhancer elements in the Roadmap Epigenomics Mapping Consortium and Encyclopedia of DNA Elements (ENCODE) revealed that human mammary epithelial cells (HMEC) have a similar epigenomic structure to alveolar epithelium, with NKX2-1 serving as a distinguishing feature of distal lung differentiation. Taken together, our results suggest that enhancer regions with dynamic transcription factor interactions are hotspots of epigenomic alteration that help to facilitate AEC differentiation.

**Author Summary:** Human health and disease states are heavily influenced by the critical cellular processes that regulate and protect our genomes. One of these safeguards is the epigenome; the coordinated set of signals overlaid on top of our DNA that controls what can happen to a given stretch DNA. Hence, epigenomic signatures play a critical role in the development and maintenance of cellular fate and function. To determine the relationship between epigenomic alterations and cellular fates of distal lung cells in humans during the process that regenerates the human lung epithelial layer after injury, we performed comprehensive genome-wide profiling of many epigenetic modifications that have roles in regulating the function of the underlying DNA. We found that changes to enhancer regions, which act to turn on associated gene expression, were the major alterations to the epigenome during distal lung differentiation, and that within those regions’ dynamic changes in transcription factor associations were occurring to facilitate this process. We then characterize what was similar and distinct to the enhancers of distal lung from among other epithelial tissues and describe a novel role for specific transcription factors in this process that previously had no known role in normal lung repair.

## Introduction

Diseases involving the distal alveolar epithelium, such as chronic obstructive pulmonary disease (COPD), idiopathic pulmonary fibrosis (IPF), and lung adenocarcinoma (LUAD), are major contributors to morbidity and mortality in the United States (1–3). While environmental factors are established contributors to the development and progression of distal lung diseases (4–6), little is understood about how the underlying epigenetic architecture of the adult lung is disrupted in these disease processes. The distal lung epithelium is comprised of two main epithelial cell types, alveolar epithelial type 1 (AT1) and type 2 (AT2) cells, each with distinct physiological roles, morphology, and molecular profiles (7). Understanding the interrelationship between these two diverse cell types and the distinct role each cell type plays in disease initiation and progression is key to developing approaches to combat peripheral lung disease.

While differences in gene expression between AT2 and AT1 cells have previously been profiled (8–12), relatively little is known about changes in the epigenetic state between these two cell types. Enhancers are epigenetic regulatory elements that control activation of gene expression and play a key role in cell type specification and regulation of disease processes (13). They are characterized by a nucleosome-depleted stretch of DNA that allows for transcription factor binding. This exposed DNA region is flanked by well-positioned nucleosomes decorated with post-translational modifications indicative of active enhancer activity. Specifically, these nucleosomes show co-occurrence of histone 3 lysine 27 acetylation (H3K27Ac) and histone 3 lysine 4 mono-methylation (H3K4me1). Open DNA regions within the center of the enhancer region can be interrogated genome-wide using Formaldehyde-Assisted Isolation of Regulatory Elements (FAIRE), followed by massive parallel sequencing (14). The open region identified by FAIRE is commonly bound by transcription factors that function to regulate downstream target gene expression levels. Often these regions are also found to be depleted of CpG methylation (15).

We set out to discover how epigenomic remodeling of AECs directs the reprogramming of AT2 into AT1 cells during AEC differentiation using a well-characterized 2-dimensional plating model derived from primary human cells. This model results in AT1-like cells, which recapitulate gene expression patterns, physiological behaviors, and morphological characteristics of AT1 cells found *in vivo (16–18)*. To do so, we performed comprehensive profiling of the epigenetic state using histone marks known to affect gene expression and regulation of genomic architecture. We identified enhancers as the epigenetic elements that influence gene expression the most during differentiation, and within them we found enrichment for high-confidence transcription factors predicted to bind to these regions. These factors could act in concert to direct AEC differentiation. We then utilized the compendium of enhancer signatures across the spectrum of human tissues to identify enhancers that were specific for human alveolar epithelial AT2 and AT1-like cells, which can be of future utility in the generation of cell-type specific models of diseases arising from the alveolar epithelium. We present herein a collection of epigenetic alterations that occur during AEC differentiation and describe their influence on coordinated gene expression patterning to direct the acquisition of an AT1-like cell fate.

## Results

### Enhancers constitute the major epigenomic alterations during AEC differentiation

We set out to determine the relationship between epigenetic alterations and AEC differentiation. To do so, we first performed comprehensive epigenomic profiling of human AEC during differentiation from AT2 to AT1-like cells. AT2 cells were extracted from explant donor lungs that had no prior evidence of chronic lung disease and allowed to differentiate into AT1-like cells *in vitro* over the course of 6 days utilizing well-established protocols (8, 19). Next, the AT2 cell population (D0), transitional AEC (D4), and AT1-like cells (D6) underwent DNA isolation for whole genome bisulfite sequencing (WGBS) (1 million cells each), chromatin fixation for ChIP-seq (5 million cells each ChIP), and corresponding RNA isolation for bulk RNA-seq (1 million cells each) to correlate altered epigenetic states with changes in gene expression from the same population of cells. Antibodies used for ChIP-seq were directed against histone modifications associated with euchromatin (H3K4me1, H3K27Ac, K3K9Ac) and facultative heterochromatin (K3K79me2/3, H3K27me3) marks, as well as the three-dimensional chromatin organizing protein, CCCTC-binding factor (CTCF). During the ChIP process, non-protein bound DNA fragments in the supernatant were collected as “free DNA” and profiled using FAIRE-seq to determine open genomic regions. Inspection of the ratio of peak enrichment to input background revealed that the ChIP-seq data were of acceptable quality for subsequent data analysis (**Figures S1-S3**). We also determined whether maximal peak occupancy was reached by subdividing ChIP-seq datasets and re-performing peak calling analysis to generate a curve for determining maximal peak occupancy (**Figures S1-S3**). Our samples had reached the plateau for number of peaks called, indicative that our sequencing depths were sufficient and had captured the vast majority of the binding sites for the given antibodies. Of note, data quality as measured by peak enrichment from Donor 1 was slightly better than Donor 2, and was therefore used as the discovery dataset, with Donor 2 used as the validation set. The genomic distribution of each epigenetic signature was then mapped back to the hg19 genome and the correlation between samples and the distribution of each mark was determined (**Figure 1A**).

**Figure 1:**
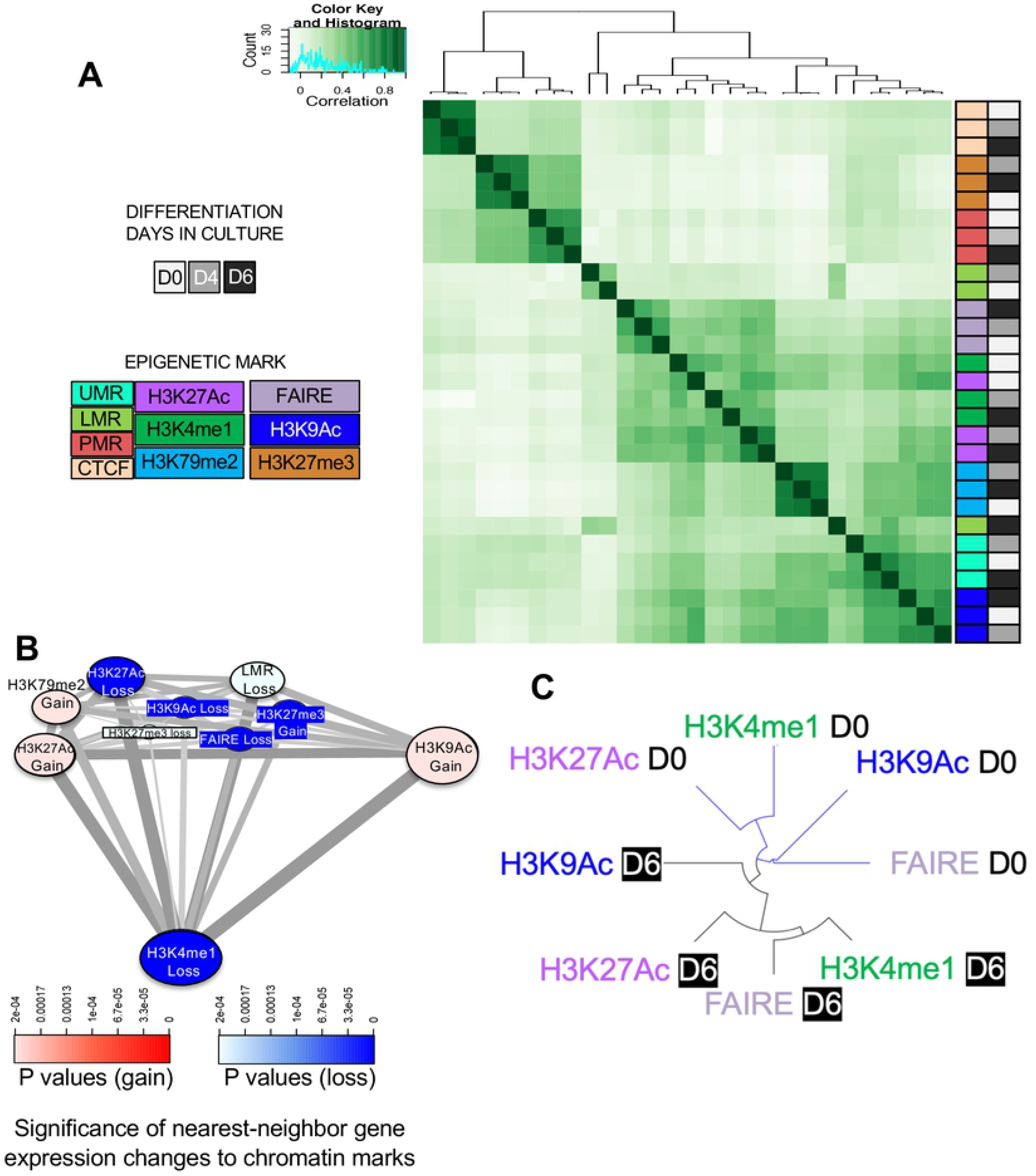
Enhancers constitute the major epigenomic alterations in AEC differentiation. A) Unsupervised hierarchical cluster analysis of R-squared correlation matrix of chromatin-mark occupancy demonstrates similarity across the major known epigenetic marks. Darker green = more highly correlated genomic distribution, white = less correlated distribution patterns across the genome. Colored annotation panels along the side of the heatmap correspond to the days in culture (greyscale) and epigenetic mark (colors) being measured. UMR = unmethylated regions, LMR = Low methylation regions, PMR = Partially methylated regions. B) PIANO diagram showing correlation between loss or gain of epigenetic mark and changes in expression of the nearest-neighbor gene. Red scale = significance of enrichment for genes with gain in expression during AEC differentiation, blue = significance of enrichment for genes with loss in expression during AEC differentiation. Increasing size of circle containing epigenetic mark = more regions associated with gene expression changes, smaller size of circle = fewer epigenetic regions associated with gene expression changes. The thickness of the grey connecting lines indicates the number of genes that are associated with both epigenetic marks, thicker = more genes associated with both marks, thinner = fewer genes. C) Dendrogram showing relationship among enhancer marks during AEC differentiation.

WGBS data underwent DNA methylation domain-calling using MethylSeekR (15), which segregates the genome into specific domains based on their level of methylation. Unmethylated regions (UMRs) have less than 10% methylation levels, extend over regions >10 kb, and have been associated with loci important for cell fate determination (20–22). Low-methylated regions (LMRs) have between 10-30% methylation levels and are associated with active enhancers (23). Partially methylated domains (PMDs) have between 30-70% overall methylation levels, tend to stretch for many kilobases (kb), and are associated with polycomb complex and facultative heterochromatin (24). The last category which is not explicitly defined by MethylSeekR are fully methylated domains (>70% methylated) which associated with constitutive heterochromatin. We integrated our WGBS domain data with the ChIP-seq data using the Diffbind package in R to calculate and visualize a correlation matrix of peak overlaps and found that partially methylated regions (PMRs) in AECs were more closely associated with the repressive chromatin mark H3K27me3 and the insulator CTCF (**Figure 1A**). CTCF acts as a long-range homodimeric insulator that regulates three-dimensional chromatin structure (25, 26). Each mark within this group clustered with itself rather than clustering together by differentiation day, indicating that these marks did not undergo major shifts during AEC differentiation.

The remaining histone chromatin marks clustered separately as active chromatin regions. UMRs clustered with the H3K9Ac mark of generalized euchromatin activation. H3K79me2, which is a mark of transcriptional elongation, segregated as their own smaller cluster within the active enhancer cluster as well. The LMR regions of D6 clustered within the active histone marks, but D0 and D4 LMR regions did not cluster with either repressive or active histone marks. The final cluster observed consisted of H3K4me1, H3K27Ac, and FAIRE signal, all marks associated with active enhancers. Interestingly, these H3K4me1 and H3K27Ac marks clustered by AEC differentiation state (i.e., days in culture) instead of by epigenetic mark, indicating that, *genome-wide*, there were substantial changes in the distribution of active enhancers as AT2 cells transition toward an AT1-like cell fate.

We have already observed that the process of AT2 to AT1-like cell differentiation alters the expression of thousands of genes (8). To further interrogate the genome-wide relationship between epigenetic state and gene expression during *in vitro* AEC differentiation, we utilized the PIANO package, which performs comparative gene-set enrichment analysis between custom datasets (27). We compared the gain or loss of each epigenetic mark profiled against changes in the HOMER-annotated nearest neighbor gene expression as a rough measure of association, with the caveat that enhancers can often target genes across great distances and in addition the rate of nearest-neighbor enhancer interaction varies across tissues and development (28) (**Figure 1B**). We observed that loss of H3K4me1, H3K27Ac, H3K9Ac, FAIRE, and gain of H3K27me3 were all highly significantly correlated to loss of nearby gene expression from differential RNA-seq analysis during differentiation (blue, all had p < 3.3×10^−5^). Loss of LMR signal was also significantly correlated to loss of gene expression, albeit to a lesser extent than the other marks (p < 2.0×10^−4^). The gain of H3K27Ac, H3K9Ac, and H3K79me3 were significantly correlated with increases in expression of nearby genes (red). None of the other epigenetic marks were significantly associated with changes in nearby gene expression. Based on these results, we decided to separate out H3K27Ac, H3K4me1, H3K9Ac, and FAIRE signals and determine the relationship among the distribution of these marks. All of the activation marks identified as associated with changes in gene expression using PIANO then underwent unsupervised clustering (**Figure 1C**). The genomic distribution of AT1-like (D6) signatures from epigenetic marks associated with enhancers were more similar to each other than to the same epigenetic mark in AT2 cells (D0). We observed that the distribution of the epigenetic signature of activation segregated based on differentiation state, rather than by the type of histone mark being profiled. Therefore, we focused on these enhancer regions critical to alterations in the epigenomic state as a means of identifying key transcriptional regulators during AEC differentiation.

### Identification of FOX family, STAT family, TEAD family, and AP1 complex members as transcription factors involved in AEC differentiation

To determine the quality of enhancer-bound chromatin mark enrichment, we plotted the overall tag density of the enhancer-associated marks FAIRE, H3K27Ac, and H3K4me1 centered on the distance from the middle of the calculated peak region (**Figure 2A**). We saw a significant enrichment of FAIRE signal at the center of each predicted enhancer, indicating that open regions were centered around transcription factor footprints as previously reported (29–31). In addition, we saw a bimodal distribution of H3K4me1 and H2K27Ac spaced ~+/−100 bp from the center of the peak, indicating nucleosomal positioning consistent with known enhancer elements as well as enrichment of enhancer-associated marks. The enrichment signal faded at ~+/−2000 bp from the center of the peak, indicating that, on average, epigenetic signals for enhancer regions extended no longer than ~4 kb. As FAIRE data most closely capture TF binding footprints in between the enhancer-decorated nucleosomes, we utilized the FAIRE data in both AT2 (D0) and AT1-like (D6) cells to look at the relative enrichment for all predicted TF motifs contained in the HOMER database (**Figure 2B**). We observed that the motifs for the TF FOS and, to a lesser extent, similar members of the AP-1 family, were the most statistically significant in the AT2 cell FAIRE regions. In contrast, we identified several TF motifs that were highly significantly enriched in AT1-like FAIRE samples, most prominently TEA domain family member - 1 (TEAD1). Notably, there were several TF motifs enriched in both cell types, such as forkhead box protein A1 (FOXA1), indicating that FOXA1 may exert its function as a pioneering TF in both cell types. To identify those motifs which demonstrated cell-type preference, we performed subtractive analysis between AT1-like and AT2 cell motif enrichment (**Figure 2C**). This demonstrated that the TEAD motifs were much more significantly enriched in the AT1-like cell motifs. The FOS motif was exclusively represented in the FAIRE open regions of AT2 cells. We observed FOS motif enrichment previously in genomic regions that had shifted histone marks from a H3K27me3 repressive state to H3K9Ac activation during AEC differentiation (8).

**Figure 2:**
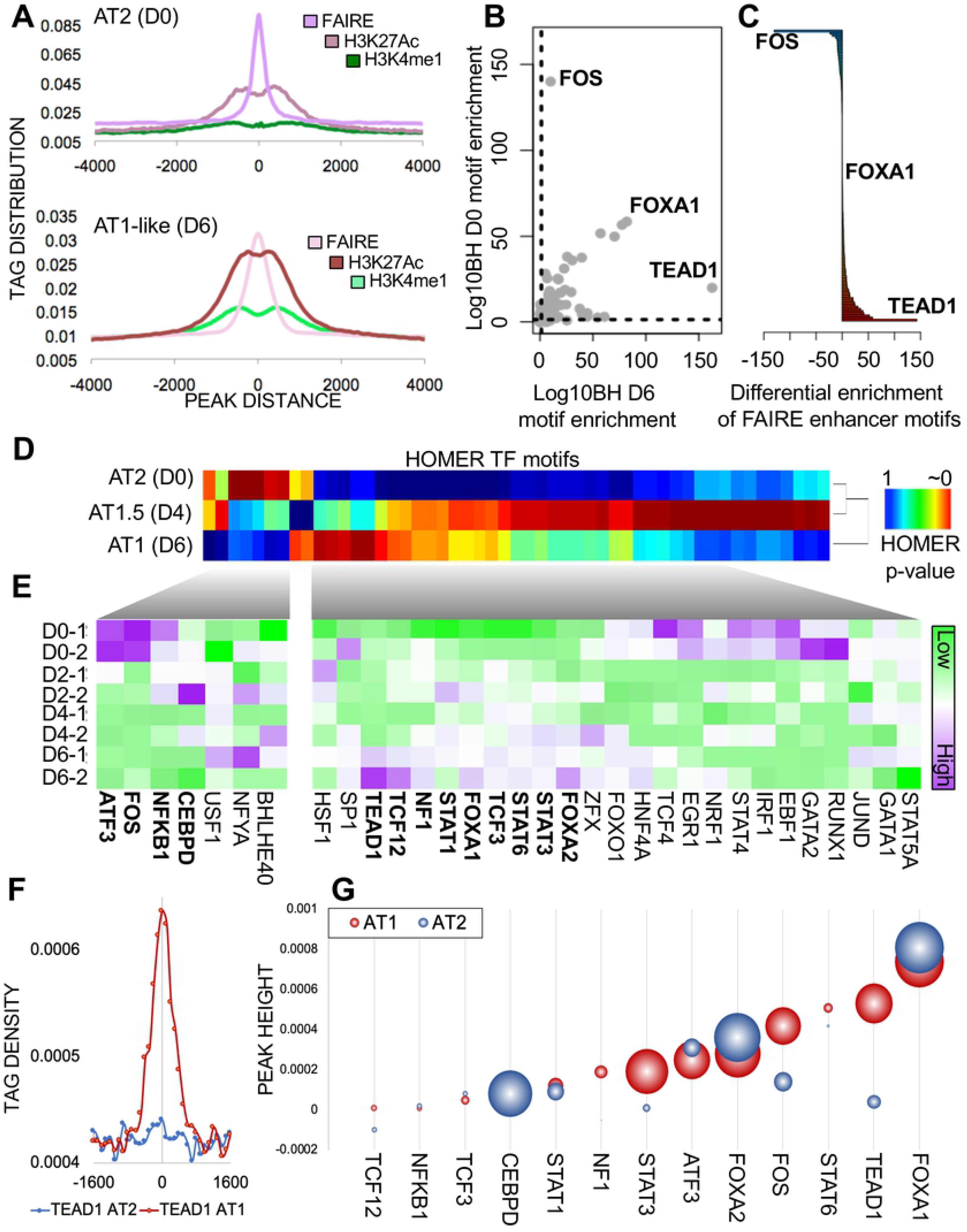
Identification of FOX family, STAT family, TEAD family, and AP1 complex members as transcription factors involved in AEC differentiation. A) Enrichment of tag density from center of the epigenetic mark for both AT2 (top) and AT1-like (bottom) cells. B) HOMER-computer enrichment of TFBS in AT1-like (X-axis, D6) or AT2 (y-axis, D0) cells. Dotted line indicates −log10BH cutoff for significance. C) Distribution of all TFBS predicted motifs available in HOMER and their enrichment in AT1-like (red) vs. AT2 (blue) cells. The BH-corrected p value for each TFBS motif was computed in each cell type and AT2 cell enrichment was subtracted from AT1-like cell enrichment. TFBS motifs were then arranged from most AT2-cell specific (top) to most AT1-like cell-specific (bottom). D) Unsupervised hierarchical clustering of TFBS enrichment in FAIRE-occupied regions. Red = Highly significantly enriched for the indicated TFBS, blue = not significantly enriched Rows are scaled based on p values of motif enrichment significance. E) Supervised clustering analysis of gene expression changes for the indicated transcription factors during AEC differentiation. Purple = High expression, green = low expression, each column color scaled by standard deviation within the row. Transcription factors bolded have loss of predicted TFBS and loss of gene expression (AT2 cells, top cluster), or gain of predicted TFBS and corresponding increase in gene expression (AT1-like cells, bottom cluster). F) Tag density of TEAD1 motif enrichment with AT2 (blue) or AT1-like (red) FAIRE peaks. Tag densities are centered on the middle of the FAIRE peak (position 0) and normalized by millions-mapped. G) The peak height (Y axis) and area under the peak (size of circle) was calculated for all AT2 (blue) and AT1-like (red) enriched TF motifs in FAIRE peaks. TFs were ranked based on AT1-like peak height (smallest to largest).

Once we had determined the statistical enrichment of TF motifs within the enhancers for each cell type, we correlated those motifs to the expression levels of their corresponding gene. Only a handful of motifs were significantly enriched in AT2 cells and not in AT1-like cells (**Figure 2D**). Comparing these predicted altered binding sites with gene expression changes throughout differentiation yielded subsets of TFs where motif enrichment in enhancers decreased along with loss of TF expression (**Figure 2E**). This set of TFs included CCAAT-enhancer binding protein delta (CEBPD), nuclear factor kappa-B (NFKB1), FOS and activating transcription factor-3 (ATF3). Consistent with these results, our previous work demonstrated a decrease of NFKB and FOS signaling during AEC differentiation (8). We also observed increased expression relative to AT2 levels, and increased motif enrichment for FOXA1, FOXA2, signal transducer and activator of transcription (STAT1/3/6), nuclear factor 1 (NF1), transcription factor 3/12 (TCF3/TCF12), and TEAD1. Previous work in our laboratory and others has demonstrated a role for FOXA1/2 and Wnt signaling in AEC differentiation (8, 32, 33).

Transcription factors are thought to create the open regions detected by FAIRE as a function of their binding. Therefore, to further refine and rank candidate TFs involved in AEC differentiation we calculated the peak height and area under the peak for each predicted TF motif in the FAIRE regions in both AT2 and AT1-like cells (see example for TEAD1 in **Figure 2F**). TFs with strong signals near the center of a FAIRE peak would fit the model that the factor is binding to the center of the FAIRE region, displacing histones and creating the FAIRE open region signal as was observed for TEAD1 in AT1 cells (red). Conversely, lack of a discernable peak near the center of the FAIRE region would argue against a functional relationship between the FAIRE open region signal and TF binding, as we observed for TEAD1 in AT2 cells (blue). We then ranked all TFs from smallest footprint to largest footprint as a measure of predictive strength of involvement (**Figure 2G**). We observed that peak height was not fully predictive of area under the peak. In addition, we observed that the HOMER calculated p value did not perfectly correlate to peak enrichment at the center of the FAIRE peak, arguing that using only the p value calculations to assign involvement of a TF may over-interpret the involvement of a given TF in the pathway being studied. Using enrichment at the center of the FAIRE peak as a metric for ranking TFs, we observed FOXA1 as the top-enriched candidate in FAIRE-marked open regions in both AT2 and AT1-like cells. In sum, we identified several TFs that are predicted to regulate enhancer dynamics and cellular phenotype during AEC differentiation. These results indicate that a single factor may not be responsible for the reprogramming of AEC, rather a network of TFs is coordinated in a temporal fashion to orchestrate gene expression changes requisite for AT1-like cell fate.

### Transcription factor interaction networks within enhancer regions shift during AEC differentiation

To confirm our previous observations that a large number of TFs were significantly enriched in AEC enhancer regions and associated with distinct sets of differentially expressed genes during differentiation, we applied knowledge from the biochemistry field about the spacing between heterodimeric TF complexes that bind site-specifically to DNA to understand how TF families were changing their associations during AEC differentiation. The majority of characterized TF heterodimeric interactions are thought to occur between binding partners that rest on DNA within 50 bp of each other, based on many decades of steric and mutational analyses (34–36). Therefore, we began by running HOMER transcription factor binding site (TFBS) prediction on AT2 and AT1-like enhancer regions. Next, we annotated where all of the top 100 significantly enriched motifs in each cell type sat within their respective enhancers. In many cases, multiple instances of a given motif were found in a given enhancer region. To reduce overrepresentation of these regions, we set a cut-off of up to 10 motif instances in a given enhancer, which encompassed over 99% of all significantly enriched TF motifs from our initial list of top 100, hereafter referred to as the “Interrogated Motif” (**Figure 3A**). Next, we ran HOMER on the 100 bp region surrounding the Interrogated Motif to determine which TF families occurred as “Associated Motifs” within that 100 bp window (blue regions, **Figure 3A**). Inclusion of the Interrogated Motif allowed for a positive control (red region, **Figure 3A**). Next, results from all 100 Interrogated motifs in AT2 cell enhancer regions (**Figure 3B**) and AT1-like cell enhancer regions (**Figure 3C**) underwent unsupervised hierarchical clustering.

**Figure 3:**
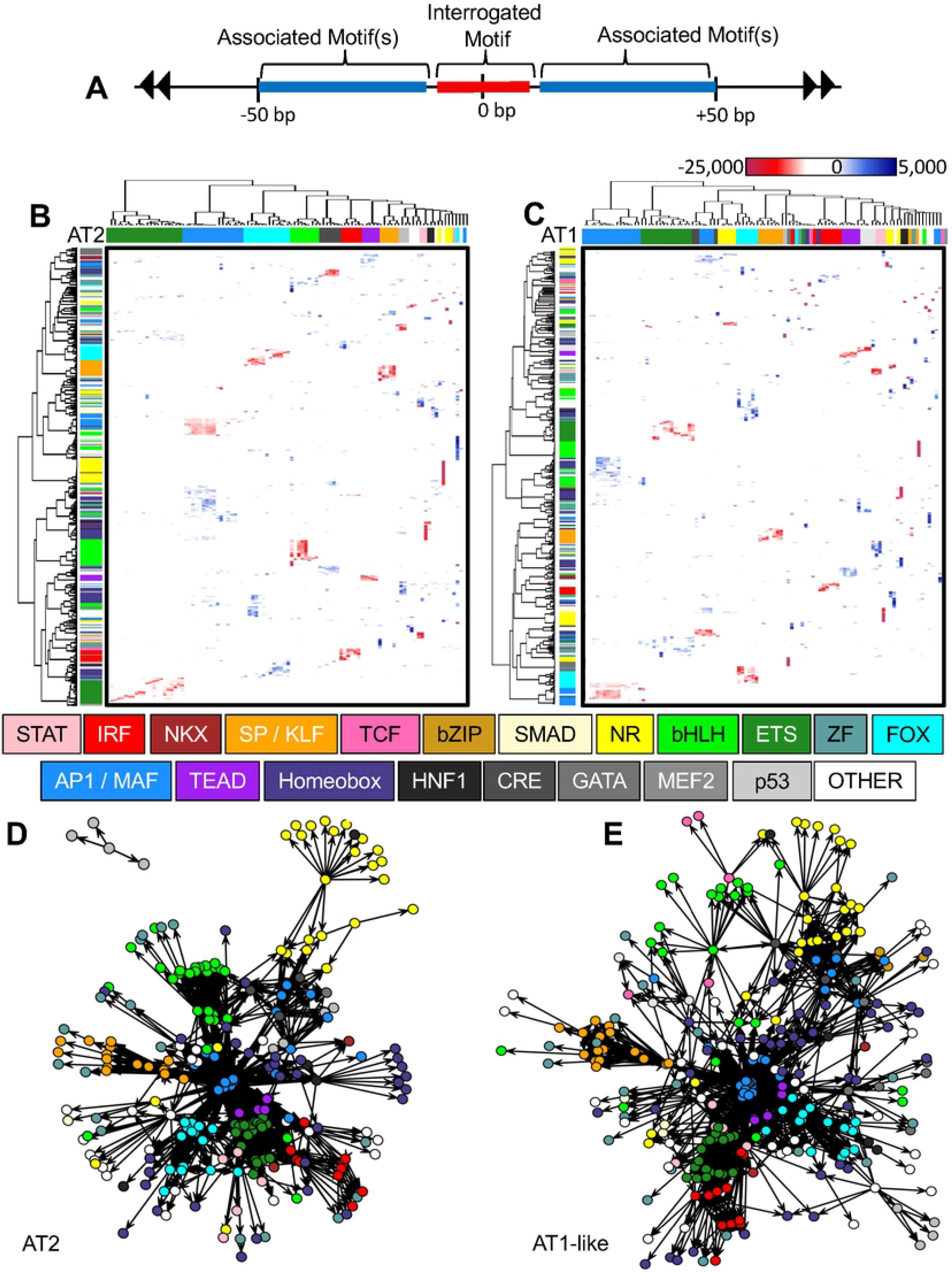
Transcription factor interaction networks within enhancer regions shift during AEC differentiation. A) Diagram of AEC enhancer regions selected for further study. Regions centered around the top 100 significantly enriched motifs within each cell type, dubbed “Interrogated Motifs”, colored red. 50 bp regions adjacent to the Interrogated Motif were subset (blue regions) to identify “Associated Motifs” that were significantly associated with the Interrogated Motifs. B-C) Unsupervised hierarchical clustering in AT2 cells (B) and AT1-like cells (C) of top 100 Interrogated Motifs (columns) and predicted Associated Motif significant interactions (rows). All HOMER TFBS were included in the analysis. Within the heatmaps, red indicates binding sequence similarity to the Interrogated Motif (positive control), blue indicates Associated Motifs had distinct core binding sequences, white indicates motif enrichment was not statistically significant. Families of TFs with similar core binding sequences were labeled with a distinct color to visually discern motif association patterns (column and row colors labels). D-E) Network analysis of AT2 (D) and AT1-like (E) enhancer TF interactions. Each circle represents a “node”, or specific TF. Families of TFs are similarly colored according to the central key within the figure. Significant association is denoted by connecting ‘edges’, ie., lines (AT2: p<e−50; AT1-like: p<e−100). Length of edge/line is not indicative of significance level, all associations above the indicated thresholds are shown.

The resultant heatmaps, showing the Interrogated Motifs as columns and Associated (secondary) Motifs as rows, showed several TF motif associations, but overall there was a large divergence in Associated Motif associations between subtypes. As expected, family members with a similar core motif sequence displayed similar enrichments for Associated Motifs, i.e., all ETS family members with the core CAGGAA sequence were predicted to have similar Associated Motif partners. This resulted in clusters of blue colored-Associated Motif families that were associated with the primary motif. Interestingly, AP1 and MAF family members share the same core TGAxxTCA sequence but differ widely in their Associated Motif association (**Figure 3B**). AP1 family members were tightly associated with ETS, FOX, and members of the bHLH family, whereas significant association of nuclear receptors (NRs) as Associated Motifs was only observed in MAF family member Interrogated Motifs. Also, in AT2 cells, TEAD family member Associated Motifs with a core sequence GGAAT were found nearby AP1 family Interrogated motifs. This is in contrast to FAIRE results, which showed enrichment for TEAD family members within FAIRE regions only within AT1 cells.

AT1-like cells showed numerous more connections between Interrogated Motif and Associated Motif families than AT2 cells. Some families, such as TCF and SMAD that were not detected in AT2 cells, were now significantly associated with multiple Interrogated Motifs. Beyond this, many families of TFs split, so that different family members, with a nearly identical core binding sequence showed drastically altered associations with Interrogated Motifs. An example of this was the Homeobox family, which was restricted to significant associations with AP1/MAF family members in AT2 cells. This was dramatically different in AT1-like cells, where homeobox cluster A (HOXA) family member Associated Motifs were significantly enriched alongside the AP1, bHLH, FOX, Zinc Finger (ZF), and TCF family members.

The high degree of interconnectivity between TF Interrogated and Associated Motifs led us to utilize a network clustering framework to visualize the degree of interaction among these families of transcription factors and how these relationships changed during differentiation. Network analysis (37, 38) was performed on the TFBS Interrogated and Associated Motifs in both AT2 cells (p<e-50, **Figure 3D**) and AT1-like cells (p<10^−100^, **Figure 3E**). Results indicated that AP1 family members formed the centralized node of each cell type’s network. Also similar was the association between IRF and ETS family members in both cell types. MAF family members cluster separately dependent on cell type and were associated with both nuclear receptors (yellow) and HNF1 family members in AT2 cells, whereas in AT1-like cells, MAF family members clustered with NRs, bZIP, Homeodomain, bHLH, and ZF family member motifs. bHLH family members (bright green) were also sequestered into their own node family in AT2 cells, with only connections to AP1 family members. In contrast, in AT1-like cells, bHLH family members were associated with TCF, nuclear receptors (NRs) (specifically retinoic acid receptor family members), CRE, Homeodomain, and zinc finger (ZF) family member motifs.

TEAD family members were associated with AP1 and ETS family members in AT2 cells. In contrast, their associations shifted in AT1-like cells to AP1, FOX, Homeodomain and ancillary other motif families. This would suggest that TEAD exchange of binding partners from ETS family to FOX/Homeodomain family members occurs without disruption of TEAD or AP1 heterodimer associations. FOX family members (cyan) were associated with primarily AP1, Homeodomain, NKX, STAT, and ETS family members in AT2 cells. In contrast, FOX family members in AT1-like cells were associated with TEAD, CRE, GATA, and MEF2C family members as well as AP1. The one exception to this pattern was the SP1/KLF family, which binds CG-rich regions that tend to form at or near transcriptional start sites regulated by CpG island promoters and remained a relatively consistent cluster. This may be reflective of our decision to not set a formal distance cut-off to exclude enhancer regions based on proximity to transcriptional start sites.

Overall, the TF network state of AT2 cells is highly ordered, with minimal connectivity of TF family nodes outside their interaction with the central AP1 TF family. In contrast, the AT1-like TFBS network displays a high degree of interconnectivity among TF family members, implicating a vast array of TFs involved in enhancer activity during alveolar differentiation. FOX family members saw a large degree of nearby Associated Motifs altered during AEC differentiation. This, combined with observations that the FOXA family members were represented in the FAIRE open chromatin regions of AT2 and AT1 cells concomitant with changes to their relative expression levels during AEC differentiation focused our attention on the interactions of this factor with known involvement in alveolar differentiation.

### FOXA1 binding in AEC is associated with TF networks in an AEC cell type-specific manner

Our work correlating epigenetic alterations with gene expression changes revealed that FOXA1 was expressed in both AT2 and AT1-like cells, was upregulated during AEC differentiation, and showed motif enrichment at the center of FAIRE-labeled open regions in both cell types. It is known that FOXA1 can translate epigenetic signatures into enhancer driven lineage-specific transcriptional patterns that coordinate cellular differentiation (39). Therefore, we decided to study the predicted binding behavior of FOXA1 in relation to other TFBS motifs within enhancers in our *in vitro* model of human AEC differentiation. The typical nucleotide spacing of TFs bound together in a heterodimeric complex is between 1-50 bp depending on the factors involved (34–36). To determine which of the identified TFs might associate with FOXA1 to maintain AT2 cellular identity or redirect FOXA1 to alternate enhancers to promote AT1-like differentiation, we gathered +/− 50 bp from the predicted FOXA1 binding site within cell-type specific enhancers and re-ran the HOMER motif analysis, excluding FOXA1 as it was a criterion for sequence selection. We found that in AT2 cells FOXA1 motifs co-occur alongside ETS family member motifs with high statistical significance (**Figure 4A**). This predicted association shifts in AT1-like cells, where TEAD family members and MEF2C are highly significantly enriched motifs alongside FOXA1. Also enriched were BAPX1 and MEF2C motifs. Consistent with these observations, we saw a decrease in ETS1 expression and increase in MEF2C during AEC differentiation, providing a possible mechanism for FOXA1 transcriptional heterodimers based on relative expression levels of cofactors. In addition, we observed enrichment of NKX2-1 and NFI in proximity of both AT2 and AT1-like FOXA1 predicted motifs.

**Figure 4:**
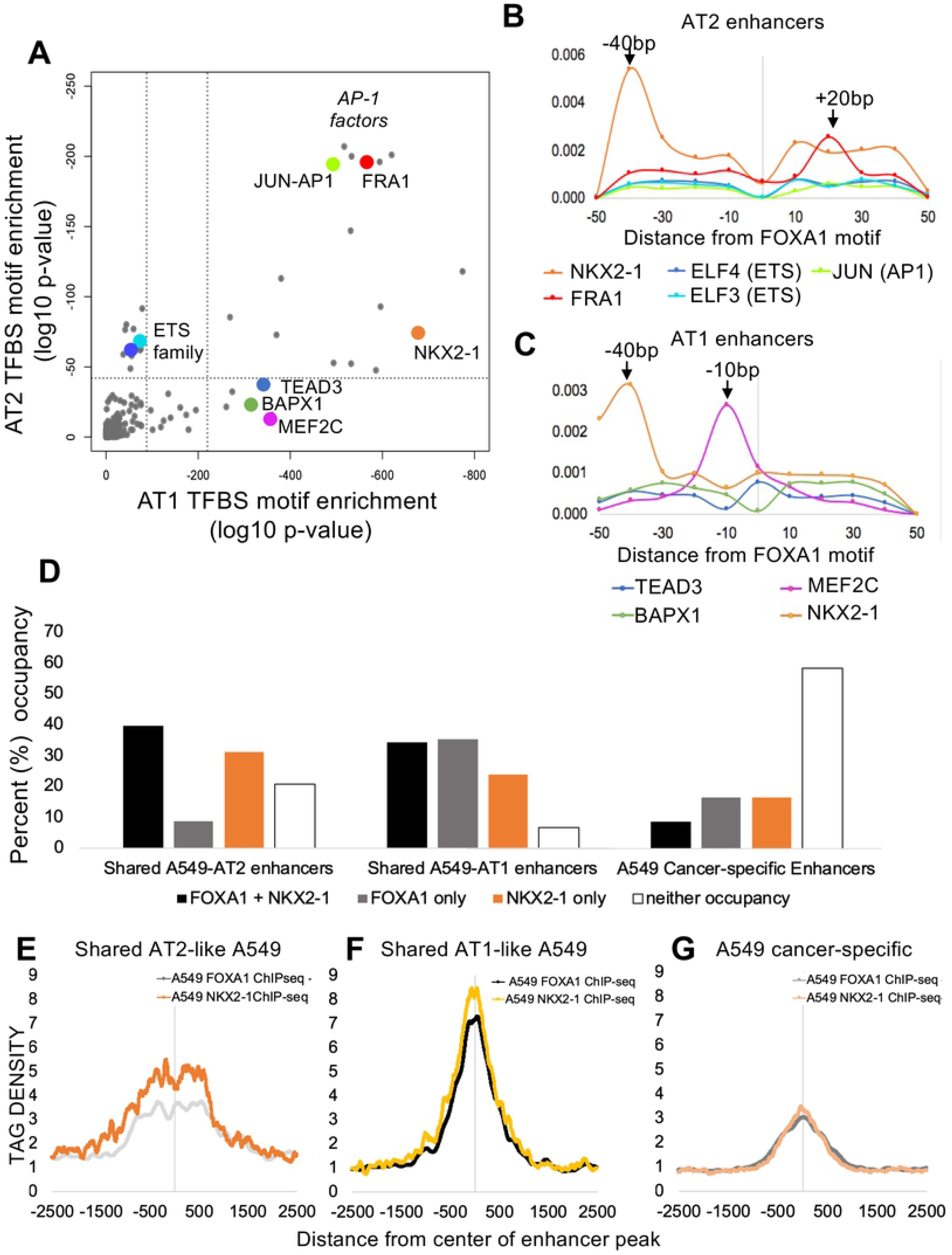
FOXA1 binding in AEC is associated with TF networks in an AEC cell-type specific manner. A) HOMER-computed TFBS enrichment for Associated Motifs surrounding the FOXA1 predicted binding motifs in AT1-like (x axis) and AT2 (y axis) cell-specific enhancers. Dotted line indicates threshold for statistical significance. B) Predicted binding site distance of the several statistically significantly enriched TFBS motifs from center of FOXA1 binding motif for AT2-cell enhancers. Zero position is the center of the FOXA1 predicted binding site. Y axis = density of predicted Associated Motif(s) at indicated bp distance from center of FOXA1 motif. C) Predicted binding site distances of the indicated TFBS motifs from center of FOXA1 Interrogated Motif for the AT1-like cell enhancers. D) Distribution of FOXA1 and NKX2-1 percent occupancy of A549 enhancers that are also present in AT2 cells, AT1-like cells, or are specific to the cancer phenotype. E-G) ChIP-seq of FOXA1 and NKX2-1 in A549 lung cancer cells. Tag density of ChIP-seq reads plotted relative to center of the enhancer peak. Tag densities between ChIP-seq runs are normalized per millions mapped. E) A549 enhancer peaks that had >50% overlap with AT2 cell enhancer regions, F) A549 enhancer peaks that had >50% overlap with AT1-like enhancer regions, G) A549 enhancers without overlap to AEC enhancers.

It is known that FOXA1 can associate with several TFs to direct cellular differentiation including NFI (40). Further, the physical interaction between FOXA1 and NKX2-1 has been observed previously in AEC (41), bolstering confidence that our analysis is identifying transcription factor complex interactions that influence epigenetic enhancer state alterations during AEC differentiation. To further characterize this relationship, we analyzed the distance from FOXA1 motifs to enriched TFBS in AT2 cells (**Figure 4B**) and AT1-like cells (**Figure 4C**). Strikingly, we observed a high degree of enrichment for NKX2-1 motifs −40 bp away from the predicted FOXA1 binding motif in both AT2 and AT1-like cells, which could indicate an interaction relationship between the two throughout AEC differentiation as previously reported (41). In addition, we observed enrichment of the FRA1/FOSL1 motif at the +20 bp position from the FOXA1 motif in AT2 cells, as well as a high level of enrichment for the MEF2C motif at the −10 bp position in AT1-like cells. In concordance with previous reports, these findings strongly indicate FOXA1 may partner with multiple transcription factors to facilitate AEC differentiation.

To determine if motif prediction was representative of actual TF factor binding patterns within enhancers, we reanalyzed publicly available ChIP-seq data that was generated in A549 cells, a cancer cell line derived from lung adenocarcinoma. AT2 cells have been thoroughly studied as a cell population that can give rise to lung adenocarcinoma (42–44). We defined enhancers in A549 cells using the same criteria as AEC, namely >50% overlap of H3K27Ac and H3K4me1 peaks. The epigenome is known to be heavily dysregulated during the carcinogenic process, so further subclassified enhancers in A549 cells by >50% peak overlap with our previously defined AEC enhancers. This resulted in enhancers of three categories: A549 enhancers that were also present in AT2 cells (1,500 regions, 5.2% of total A549 enhancers); A549 enhancers that were also present in AT1-like cells (9,678 regions, 32.8% of A549 enhancers); and A549 enhancers that were uniquely present in the cancerous cell line (18,303 regions, 62% of A549 enhancers) and may therefore may represent dysregulated enhancer activity. To determine TF occupancy within these categories of enhancers, we reanalyzed ChIP-seq data for endogenous FOXA1 originally generated by the ENCODE Consortium (45), and a separate study that determined occupancy for ectopically expressed NKX2-1 in A549 cells (46). Unfortunately, publicly available MEF2C and Fra1/FOSL1 ChIP-seq datasets were not available in lung-derived cell lines.

Overall, only 13.9% of A549 cell enhancers exhibited co-occupancy of FOXA1 and NKX2-1 by ChIP-seq. However, we observed differences in co-occurrence from this average depending on whether the A549 enhancer was categorized as ‘shared with AT2’, ‘shared with AT1’, or ‘A549 cancer-specific’. For shared A549-AT2 enhancers, 39.6% had co-occupancy of NKX2-1 and FOXA1 (**Figure 4D**). Similarly, NKX2-1 and FOXA1 peaks were co-occurrent in 34.3% of A549-AT1 enhancers. In contrast, A549 cancer-specific enhancers contained considerably fewer instances of FOXA1 and NKX2-1 peak co-occurence (8.6%). Together this indicated that co-occupancy of FOXA1 and NKX2-1 within “normal” AEC enhancers occurred approximately three times more often than within A549 cancer-specific enhancers. Indeed, almost 60% of cancer-specific A549 enhancers lacked any binding for FOXA1 or NKX2-1 (**Figure 4D**), suggesting that the colocalization of FOXA1 and NKX2-1 observed in A549 cells is primarily driven by enhancers preserved in normal tissues

To determine the relative positioning of FOXA1 and NKX2-1 in the cell type-specific subsets of enhancers, we extracted sequence alignment map (SAM)-level data and used HOMER to generate Tag densities at the cell type-specific peak regions. In AT2 cell-type enhancers that are also present in A549 cells, FOXA1 and NKX2-1 exhibited enrichment was spread across the central 500 bp of the enhancer peaks (**Figure 4E**). In contrast, AT1-like cell enhancers also present in A549 cells showed a high degree of enrichment for both factors toward the central 100 bp of the enhancer peaks (**Figure 4F**). In cancer-specific enhancers in A549 cells, there was far less enrichment for both NKX2-1 and FOXA1, with no obvious differences in TF position relative to the center of the peak (**Figure 4G**). Intriguingly, the NKX2-1 and FOXA1 datasets both exhibited a dip at the exact center of the peak for AEC-shared enhancers, which may be due to the presence of another factor. To investigate what factor might be bound there, we extracted the central 100 bp from those AT1-like enhancers shared with A549 cells that also had co-occupied by NKX2-1 and FOXA1. JunB (p=3.2×10^−16^) and MEF2C (p=1.4×10^−8^) were the predicted factors to bind this center-of-the-peak region. This could indicate that FOXA1 and NKX2-1 operate in a trimeric complex with either MEF2C or AP1/JunB family members.

### Identification of NKX2-1 and MEF2C as FOXA1-associated TFs that specify lung epithelium differentiation

Once we had characterized the relationships between TFBS motif enrichment and epigenetic state alterations during AEC differentiation, we sought to determine if the predicted interactions were unique to lung differentiation or a common phenomenon shared among other cell lineages. To investigate this, we utilized publicly available high-quality ChIP-seq datasets from normal tissues profiled by the ROADMAP epigenomics project (76 samples) and ENCODE (6 samples) (45, 47). To define what an enhancer was across multiple tissue types, we used the criterion that each cell type needed to have high-quality ChIP-seq data for H3K27Ac and H3K4me1. The H3K27Ac peaks in each cell type were then filtered to include only those that had >50% overlap with H3K4me1 peaks in the same cell type.

Diffbind analysis showed clustering of embryonic stem (ES)/induced pluripotent stem (iPS) cells as distinct from all other cell types (**Figure 5A**). Hematopoietic lineages also clustered separately from other tissues (including purified blood cell types, thymus and spleen). Interestingly, epithelial (light blue) and mesenchymal (light green) cell types were more similar to each other than all other cell types examined, with AECs closely related to the epithelial datasets present, which were human mammary epithelial cells (HMEC) and foreskin. We saw slight variation in the cell types most associated with AEC when clustered by H3K27Ac or H3K4me1 marks individually (**Figure S4**); however, breast epithelium was consistently one of the most closely associated tissue by epigenetic signatures that were available from ROADMAP and ENCODE.

**Figure 5:**
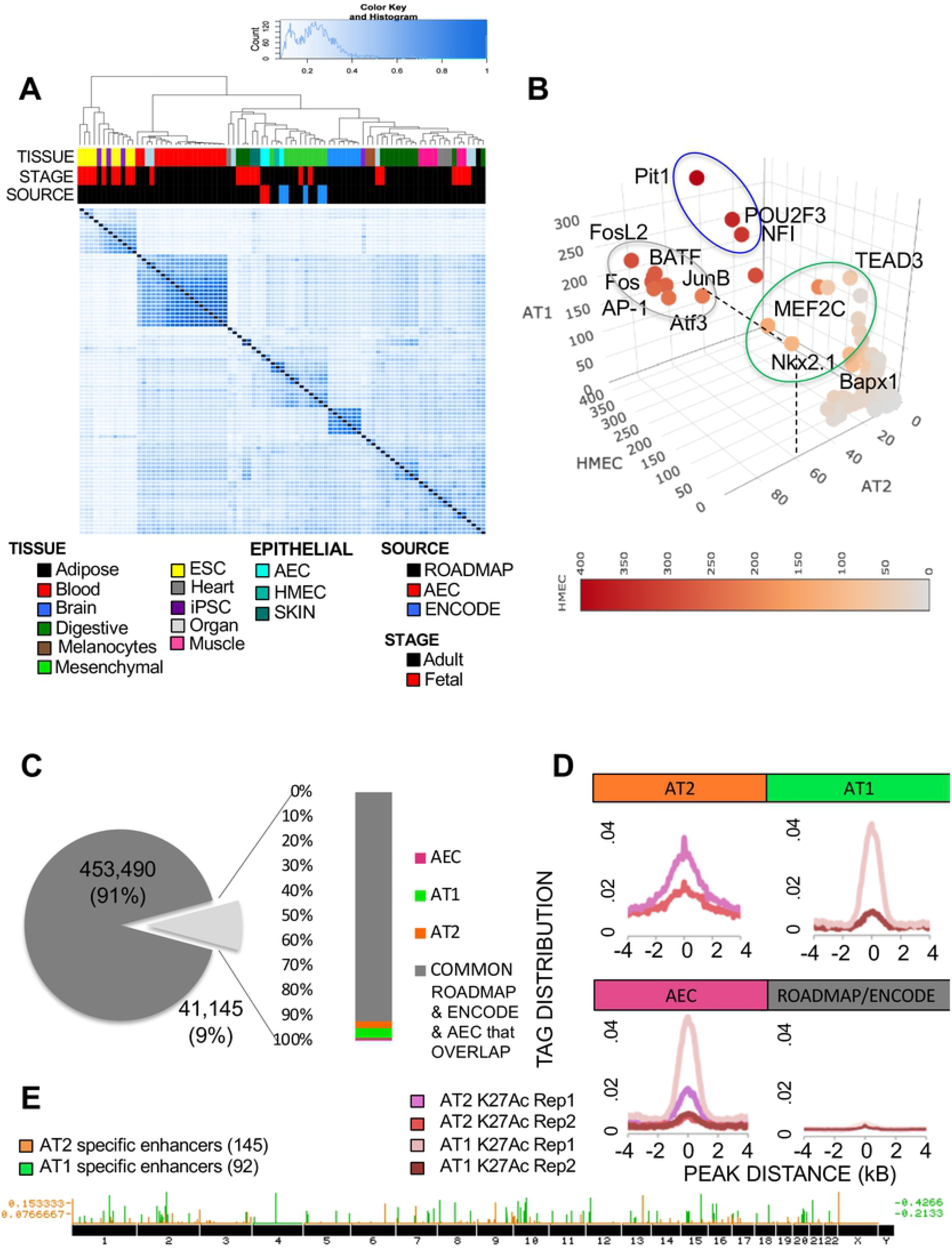
Comparative analysis of AEC enhancers with ROADMAP and ENCODE enhance-ome reveals TFBS motifs with specific enrichment in AT2 and AT1-like cell types reveals 237 alveolar epithelial-specific enhancers. A) Diffbind plotting of similarity between enhancer regions. Tissue = ROADMAP or ENCODE indicated cell type. Stage = age of donor, subdivided into pre- and post-natal. Source = origin of the data used in the analysis. B) Three-dimensional scatterplot of enrichment for each of the TFBS present in HOMER in AT2, AT1-like, and HMEC enhancers. Red scale coloring indicates level of enrichment in HMEC. Grey circle indicates TFBS motifs enriched in all 3 cell types. Blue circle indicates TFBS enriched in HMEC and AT1-like cells, green circle indicates TFBS motifs enriched specifically in AT2 and AT1-like cells. C) Pie chart indicating similarity between ROADMAP/ENCODE enhancers and AEC-identified enhancers. AEC = regions labeled enhancers in both AT2 and AT1-like cells but not in any ROADMAP or ENCODE dataset. D) Histogram indicating tag-density of enhancer-specific enrichment for H3K27Ac between biological replicates. E) Distribution of AT2 and AT1-like cell-specific enhancer regions genome-wide.

Breast and lung both undergo branching morphogenesis, and FOXA1 has a demonstrated role for both tissues in this process (48, 49). Therefore, we wanted to determine if the transcriptional co-network of TFs associated with FOXA1 was common to both or if instead, FOXA1 transcriptional networks varied between these tissues. To do so, we evaluated motif enrichment within 50 bp of FOXA1 predicted sites in primary human mammary epithelial cell (HMEC) enhancers, repeating the process that was done in Figure 3A. In total, 45% of AEC enhancer regions overlapped with HMEC enhancers, suggesting that although HMEC was the most closely related cell type studied, there was still considerable variation between their epigenetic states. 53% of all enhancer peaks in HMEC contained the predicted FOXA1 binding motif. Motif enrichment analysis was then re-run on the 100 bp surrounding the predicted FOXA1 binding sites in HMEC enhancers. Because enhancer regions were selected based on the presence of FOXA1, FOX family motifs with similar sequence to FOXA1 were eliminated from the subsequent analysis. A three-dimensional scatter diagram of enrichment measurements for all available TFBS motifs in AT2, AT1-like, and HMEC enhancer regions (**Figure 5B**). Enrichment for FOS/JUN (AP-1) motifs was observed in proximity to FOXA1 in all three tested cell types (grey circle), indicating that partnering between FOXA1 and FOS/JUN factors may play a conserved role in epithelial cell types. A separate cluster of TFBS motifs enriched in AT1-like cells and HMEC also emerged (blue circle), which included PIT1, POU2F3, and NF1. NF1 is a known binding partner of FOXA1, whereas POU2F3 and PIT1 are involved in cellular fate determinations. This could be reflective of the role these factors play in cellular differentiation (50–52). Lastly, a separate cluster of TFBS enriched in AT2 and AT1 but not HMEC was observed (green circle). This included NKX2-1, a known lung-specific lineage factor, as well as TEAD family members and MEF2C. Therefore, we have identified a high confidence set of transcription factors that appear to act in concert to coordinate AEC differentiation *in vitro* and distinguish between lung and breast enhancer identity.

### Identification of AT2 and AT1-like enhancers unique to AEC from the known compendium of human enhancers

To determine how these transcription factor coregulatory networks described above work in concert to specifically activate cell type specific enhancers, we first identified enhancer regions that were present only within AEC. The considerable variation in enhancer location across all tissues present in ROADMAP/ENCODE and the observation that enhancer regions best recapitulated the epigenetic signature of differentiating AECs gave rise to the idea that we could utilize publicly available datasets on enhancer locations to define AEC cell-specific enhancer signatures for both AT2 and AT1-like cells. To do so, the entire complement of ROADMAP (47) and ENCODE (45) enhancers for the 82 normal cell types across many organ types was merged to create one master list containing all regions within the human genome identified as enhancers, which we will refer to as the “enhance-ome”. Cancer-derived enhancer signatures were omitted due to their potential perturbation by the carcinogenic process. The locations of AT2 and AT1 cell enhancer regions were then compared to the enhance-ome. AECs had 41,145 active enhancers at 9% of all identified normal enhance-ome regions (**Figure 5C**). Of those 41,145 sites in AECs, 92% were also considered enhancers in ROADMAP and ENCODE data sets, providing us with a high level of confidence that our AEC-defined enhancers were consistent with observations from other sources. Within the enhancers present in AEC cells but not in ROADMAP or ENCODE, 295 enhancer regions were active in both AT2 and AT1-like cells (termed AEC), 1277 enhancer regions were only active in AT2 cells (ie., not present in AT1-like, ROADMAP or ENCODE), and 1706 enhancer regions were only present in AT1-like cells.

To validate these regions as either AT2 or AT1 cell-specific enhancers, we utilized the biological replicate ChIP-seq data from Donor 2. H3K27Ac peak enrichment was centered similarly between Donor 1 and Donor 2 in both AT2 and AT1-like samples (**Figure 5D**); however, the overall enrichment was lower for the biological replicate from Donor 2. Subsetting the AT2 cell-specific and AT1-like cell-specific peaks from Donor1 to overlap with peaks called from Donor2 resulted in identification of 145 AT2 cell-specific and 92 AT1 cell-specific high-confidence enhancers (**Figure 5E**).

### MEF2C:FOXA1:NKX2-1 transcription factor heterotrimeric complexes are enriched in AT1 cell type specific enhancers

Although we identified transcription factor co-regulatory networks as well as AEC cell-type specific enhancer regions, the influence of the FOXA1-associated TFBS on cell type-specific enhancer regions remained unanswered. To address this, we analyzed the distribution of TFBS motifs within the AT2 and AT1-like cell-type specific enhancers and found that all of the AT1-like cell-type specific enhancers had motifs for at least one of the TFs that were identified as associated with FOXA1 in AT1-like cells. The majority of AT1-like cell-specific enhancers had predicted motifs for all three TFs: FOXA1, NKX2-1, and MEF2C (**Figure 6A**). Many of the AT1-like cell-specific enhancers had predicted motif distributions consistent with the TF spacing we observed previously, consistent with what is known about the interaction of these TFs in the literature (**Figure 6B**). To determine the relationship between FOXA1 and NKX2-1 positioning in AT1-like cell type-specific enhancers, relative FOXA1 and NKX2-1 ChIP-seq tag density enrichment was plotted across the 92 AT1-like cell-specific enhancers from the publicly available A549 datasets (**Figure 6C**). We observed that staggered spacing between FOXA1 and NKX2-1 peak summits offset was larger in the ChIP-seq data than from motif prediction (190 bp in ChIP-seq vs 40 bp in motif prediction), which may be due to a loss of resolution in the ChIP-seq due to fragmentation size of the ChIP libraries. Enhanced association between FOXA1, NKX2-1, and MEF2C may be explained by significant increases in expression during alveolar differentiation (**Figure 6D**). Our results suggest the association of FOXA1, NKX2-1,and MEF2C may act in a cooperative heterotrimeric TF complex which binds to AT1-like enhancers as part of a coordinated effort to differentiate the alveolar epithelium, which is reflected in concomitant alterations to the epigenetic state to mediate cellular fate determination.

**Figure 6:**
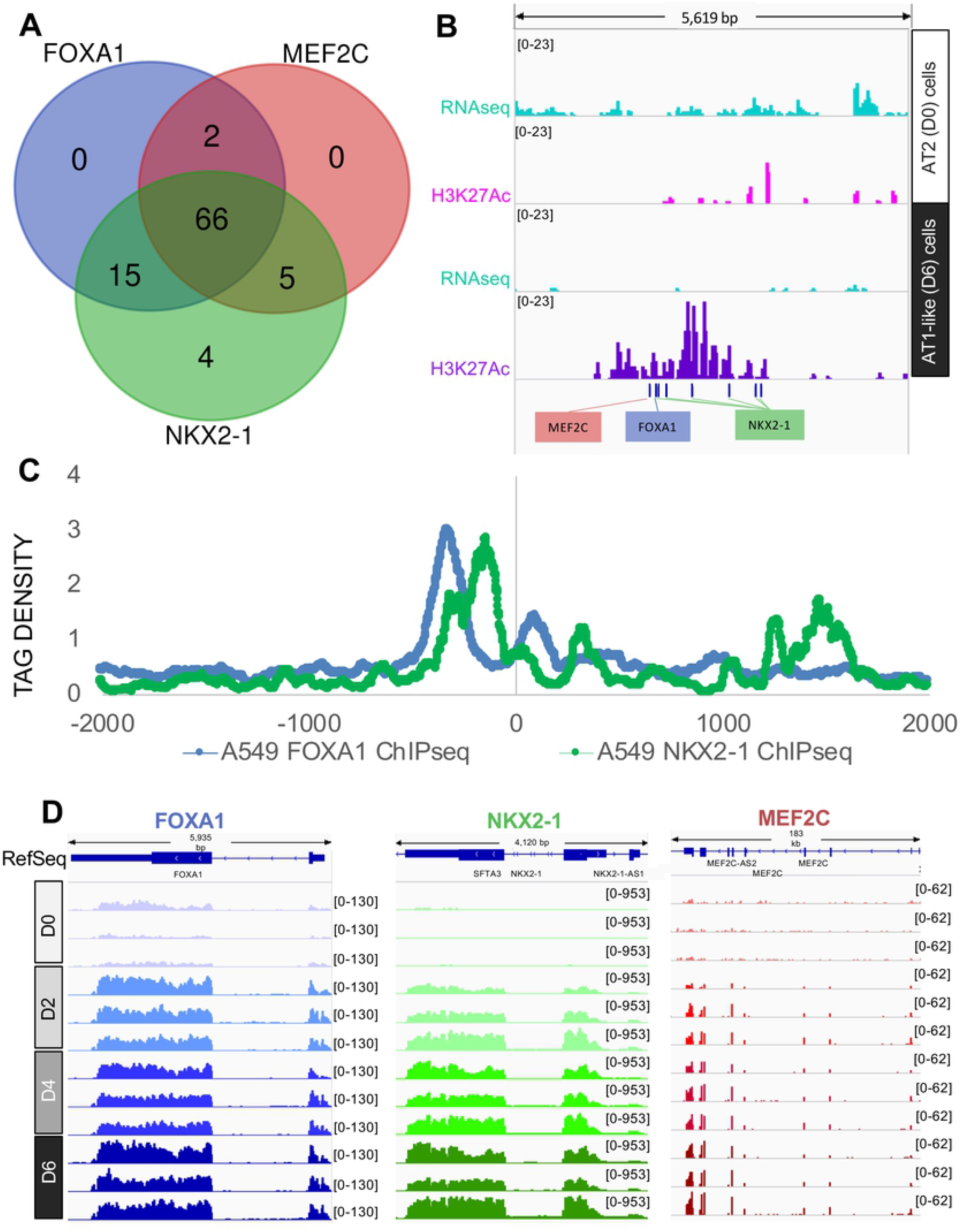
MEF2C:FOXA1:NKX2-1 transcription factor heterotrimeric complexes are enriched in AT1-like cell type specific enhancers. A) Presence of predicted NKX2-1, MEF2C, and FOXA1 binding sites within the 92 AT1-like cell-specific enhancer regions. B) Example of one specific locus with an AT1-like cell type specific enhancer and the relative positioning of predicted FOXA1, NKX2-1, and MEF2C TFBS. C) A549 cell line ChIP-seq of FOXA1 (blue) and NKX2-1 (green) distribution in AT1-like cell-specific enhancers. Tag Densities are centered on the middle of the cell type specific enhancer peak. D) IGV display of FOXA1 (blue), NKX2-1 (green), and MEF2C (red) expression levels during AEC differentiation. D0 = Day 0 (AT2), D2 = Day 2 (AT1.5, intermediate), D4 = D4 (AT1.5, intermediate), D6 = Day 6 (AT1-like).

## Conclusions

We set out to characterize the extent of epigenomic alterations that occur during AEC differentiation and how they influence cellular identity. We found that the enhancer-associated epigenetic signatures of FAIRE open regions, H3K27Ac peaks, and H3K4me1 peaks were most closely associated with changes in gene expression during AEC differentiation. Exploring this linkage further, we found that the composition of predicted TFBS motifs changed dramatically during *in vitro* AT2 to AT1-like cell differentiation, with some TFs (e.g., FOS, ETS1, and NKFB1) enriched in AT2 cell maintenance losing expression and simultaneously having decreased predicted binding to enhancer regions in AT1-like cells. Others, such as MEF2C and TEAD family members, increased in expression and had corresponding increases in predicted TF binding when transitioning to an AT1-like cell fate.

We also found that the transcription factor FOXA1, known to regulate branching morphogenesis of the lung and AT2 cell maintenance, may play a critical role in human AT2 to AT1 cell differentiation by partnering with the lung-specific transcription factor NKX2-1. This may be accomplished by switching TF heterotrimeric complex members during differentiation, as we observed differential enrichment for FRA1/FOSL1 and MEF2C in AT2 and AT1-like cells, respectively. This heterotrimeric complex member switching could facilitate alternate enhancer target localization or alter the function of the complex. Interestingly, the previously reported NKX2-1:FOXA1 interaction at the SFTPC promoter was deemed inhibitory (41), whereas we observe these predicted interactions in regions bearing epigenetic marks characteristic of “active enhancers” associated with transcriptional activation. Additionally, it has been previously reported that loss of NKX2-1 can direct the FOXA1/FOXA2 TF axis to alter cell fate from lung to stomach phenotypes (53), specifically for AT1 cells (54). Our analysis provides a basis for connecting these disparate lines of evidence. Namely, that beyond the known role of NKX2-1 in establishment of the lung endodermal lineage from thyroid (55, 56), and the role of FOXA1 in lung branching morphogenesis (49), the FOXA1:NKX2-1 interaction may be pivotal in regulation of epigenomic fate during AEC differentiation.

Interestingly, PIANO analysis revealed that FAIRE gain was also significantly associated with downregulation of associated gene expression, perhaps as a result of repressor factor occupancy at sites outside of active enhancer regions. Indeed, analysis of overall FAIRE binding sites failed to identify several FOXA1-associated TFs when enhancer regions were instead selected based on co-occupancy of H3K27Ac/H3K4me1 marks. While determining the precise functional role for any of the TFs we have uncovered in this study will require further *in vitro* characterization, we provide here a compendium of highest-priority TF candidates that recapitulate on a genome-wide scale our previous *in vitro* findings that were determined at individual loci.

We also investigated conservation and uniqueness of AT2 and AT1-like TF co-regulatory networks by examining the significance of individual TFBS motif enrichment in the most closely related cell type profiled by the ROADMAP and ENCODE databases, that of human mammary epithelial cells (HMEC). We discovered a subset of FOXA1-associated TFs common to all three cell types, and enrichment for FRA1/FOSL1 at +20 bp downstream of the FOXA1 motif. Therefore, FRA1/FOSL1 interaction with FOXA1 may play a critical role in multiple organs. Follow-up work in mouse models will be important to determine if conditional knockout of FRA1/FOSL1 is able to recapitulate the deleterious effects on branching morphogenesis seen in FOXA1/FOXA2 double knockout mice (49).

It should be noted that members within the same family of transcription factors often have nearly identical TFBS motifs. Throughout the paper, we refer to specific TFBS as enriched based HOMER motif predictions; however, in the absence of confirmatory ChIP-seq (DNA occupancy) and RNA-seq (expression) data, these motifs could be bound by any one or multiple TF family members with similar DNA binding preferences. For example, the FOXA motif is nearly identical for all three FOXA family members (FOXA1, FOXA2, and FOXA3), limiting occupancy predictions based purely on motif enrichment to a family of related TFs rather than implicating a specific TF. Complicating the interpretation of FOXA motif enrichment is their known compensatory roles in branching morphogenesis of the lung and alveolar epithelial differentiation (49).

We also identified a high-confidence set of AEC cell type-specific enhancers that were present in biological replicates of AT2 or AT1-like cells, but not present in other ROADMAP or ENCODE normal tissue databases. We found that key transcription factor coregulatory network partners identified in our genome-wide analysis were also present at highly selective AT1-like specific enhancer sites. This would support the notion that there is no one specific TF that drives AT1-like enhancer activation but is instead the result of combinatorial TF activity that is acting broadly across the genome and a very small percentage of these sites happen to occur distinctly in AT1-like cells.

In summary, we have identified epigenetic signatures characteristic of primary human alveolar epithelium and have elucidated mechanistic insights into how these shift in an *in vitro* model of primary AT2 to AT1-like cellular differentiation. These epigenetic signatures are being made publicly available to further understanding of alveolar epithelial cell differentiation process with particular emphasis on how epigenetic signatures dictate the coordinated pathways that result in altered cellular fate (7, 57). AEC differentiation *in vitro* from purified adult human, rat, and mouse AT2 cells is considered a model of wound healing (58, 59), as adult AT1 cells must be replenished after exposure to and damage from a slew of particulate and chemical insults present in the air we breathe (60). The ability of the TFs we have identified to facilitate this process may be affected by these environmental insults, leading to disrupted AEC differentiation and wound healing. This can in turn manifest as diseases of the distal alveolar epithelium, such as idiopathic pulmonary fibrosis, chronic obstructive pulmonary disorders, and lung adenocarcinoma. Importantly, many of the transcription factors we have identified in this study have known roles in these disease processes. Specifically, FOXA1 plays a significant role in non-small cell lung cancer (61), MEF2 family members have a role in lung carcinoma (62, 63), TEAD family members have known roles in carcinogenesis of epithelial tissues (64), and NKX2-1 has a long history of involvement in lung cancer, COPD and IPF (65–69). Understanding the relationship between disruption of the epigenetic state during AEC differentiation and the development of lung diseases could open up an entirely new avenue of therapeutic options for these often-fatal diseases.

## Declarations

### Ethics Approval and Consent to Participate

Remnant human transplant lung was obtained from donors in compliance with protocols for the use of human source material in research (HS-07-00660).

## Data Availability Statement

All newly generated datasets used in this study are deposited in the public GEO database (GSE150527). ENCODE data, including cell lines and normal tissues data as well as ethanol treated FOXA1 ChIP-seq in A549 cells can be publicly accessed from the UCSC genome browser (https://www.genome.ucsc.edu/ENCODE). ROADMAP epigenomics consortium data can be publicly accessed from wwww.roadmapepigenomics.org. ChIP-seq of lentiviral-introduced NKX2-1 in A549 cells was previously published (67). All analysis software used to generate used in these analyses are publicly available. All materials are commercially available from the vendors listed in the Materials & Methods section.

## Acknowledgements

The authors would like to acknowledge the efforts of the USC Epigenome Core and its former director, Charles Nicolet, for the generation of NGS data contained within this study. We would also like to thank Dr. Peggy Farnham for her expertise and recommendations as to which ChIP-seq protocols and antibodies would work best on primary cells. We also thank J. Alvarez, M. Flores and H-J. Wang for assistance with AT2 cell preparation and AT1-like cell differentiation as well as RNA isolation. Funding for this project was provided by R01 HL114094-01 to IAO and ZB, R35 HL135747 to ZB, as well as the Hastings Foundation. Funding for the Epigenome Core was supplemented by the Norris Comprehensive Cancer Center core grant P30 CA014089-43. ZB was the Ralph Edgington Chair in Medicine at USC throughout the development of this manuscript. CNM was supported in part by an American Cancer Society – Research Scholar Grant [RSG-20-135-01-RMC]. The authors declare no conflicts of interest.

## Author Contributions

CNM, BZ, KS, IAO, and ZB conceptualized the experiments and analysis. BZ, TRS, YL, JL, MER, ET, and CNM performed sample and data collection. CNM, BZ, DM, LM, YW, ET, EAM, ALR, IAO, ZB wrote and edited the paper. CNM, DM, SKL performed the analysis. TRS, LM, YW performed validation and functional analysis of results. CNM, IAO, and ZB supervised the project. Funding acquisition was provided by ZB, IAO and CNM.

## Materials & Methods

### Isolation and culture of human alveolar epithelial cells

Donor lungs were processed within 3 days of death. Lung tissue processing protocol was modified from (8, 70). In brief, AT2 cells were isolated from cadaveric human lungs that were declined by the Northern California Transplant Donor Network. Donor 1 was a 62-year-old Caucasian male. Donor 2 was a 25-year-old Caucasian male. Neither died from lung-related injury or complications. We selected the lobe of the lung that had no obvious consolidation or hemorrhage by gross inspection. Previous studies indicate that these lungs are generally in a relatively normal condition physiologically and pathologically. Cells were isolated after the lungs had been preserved for 4–8 h at 4°C. The pulmonary artery was perfused with a 37°C PBS solution, and the distal air spaces were lavaged with warmed Ca^2+^- and Mg^2+^-free PBS solution (0.5 mM EGTA and 0.5 mM EDTA) 10 times. Next, 13 U/ml elastase in Ca^2+^- and Mg^2+^-free HBSS were instilled into the distal air spaces through segmental bronchial intubation. After digestion for 45 min, the lung was minced finely in the presence of fetal bovine serum (FBS) and DNase (500 μg/ml). The cell-rich fraction was filtered by sequential filtration through one layer of sterile gauze, two layers of gauze, and 150-μm and 30-μm nylon mesh. The solution was then layered onto a discontinuous Percoll density gradient 1.04–1.09 g/ml solution and centrifuged at 400 *g* (1,500 rpm) for 20 min. The upper band containing a mixture of AT2 cells and alveolar macrophages was collected and centrifuged at 800 rpm for 10 min. The cell pellet was washed and resuspended in Ca^2+^- and Mg^2+^-free PBS containing 5% FBS. The remaining cell suspension was incubated in human IgG-coated tissue culture-treated Petri dishes in a humidified incubator (5% CO_2_, 37°C) for 90 min. Unattached cells were collected and counted. The cells were then incubated with FcR blocking (Miltenyi Biotec #130-059-901), CD326 (EpCAM) beads (Miltenyi Biotec #130-061-101) and rotate 10’@4°C followed by 20’@ RT. Collect the cells using LS columns (Miltenyi Biotec #130-042-401) on the magnetic stand. Cell viability was assessed by the trypan-blue exclusion method. AT2 cell purity was determined by NKX2-1 cytospin staining (1:100, Leica Biosystems, Cat # NCL-1-TTF1). Donor1 was 83% positive, Donor 2 was 96% positive. AT2 cells were then plated at 4×10^6^ cells per well in 6-well Corning plates and incubated at 37°C in 50:50 DMEM high glucose:DME-F12 media supplemented with 10% fetal bovine serum (FBS). 1 million cells were used for DNA and RNA extraction, 2-5 million cells were used for chromatin immunoprecipitation (ChIP-seq) of histone marks, and 10 million cells were used for CTCF ChIP-seq.

#### Extraction and processing of RNA for bulk RNA-seq and DNA for whole-genome bisulfite sequencing

1 μg of total RNA was isolated from the indicated alveolar epithelial cells using the Illustra TriplePrep Kit (GE LifeSciences, Piscataway, NJ). RNA underwent library preparation and sequencing on the IlluminaHiSeq2000 at the USC Epigenome Core. Briefly, total cell RNA was DNase I digested and then subjected to ribosomal RNA depletion with the Ribominus™ Eukaryote v2 kit (Life Technologies, # A15020, Grand Island, NY). Libraries were constructed with the TruSeq RNA Sample Prep Kit v2 (Illumina # RS-122-2001) and underwent Illumina HiSeq 2000 paired-end sequencing (2 × 50 bp) according to the manufacturer’s instructions as previously reported (31, 71). Resultant 50 bp paired end FASTQ files were trimmed to remove adapters and realigned to the hg19 genome using Bowtie 2 (72). Mapped reads were then assembled into transcripts using TopHat v2.0.12 (73). Resultant reads per kilobase of gene per millions mapped (RPKMs) were used for downstream analysis included herein. For whole genome bisulfite sequencing (WGBS), DNA was isolated and library preparation was performed as at the USC Epigenome Core. In brief, libraries were plated using the Illumina cBot and run on the Hi-Seq 2000 according to manufacturer’s instructions using HSCS v 1.5.15.1. Bisulfite-treated DNA underwent Paired End 100 cycling; Image analysis and base calling were carried out using RTA 1.13.48.0, deconvolution and fastq file generation was carried out using CASAVA_v1.7.1a5. Alignment to the genome was carried out using bsmap V 2.5 (74). Aligned .bam files were visualized using IGViewer V2.3.40 (Broad Institute, Cambridge MA). Reads were then aligned to the hg19 bisulfite genome and CpG methylation levels and SNPs were determined genome-wide using BisSNP (75). Methylation domains for each time point during differentiation were calculated using MethylSeekR (15).

#### Generation of ChIP-seq and FAIRE from primary human epithelial cells

Chromatin immunoprecipitation (ChIP) was performed using antibodies (Abs) against H3K27Ac (Cat # 39133, Active Motif, Carlsbad CA), H3K4me1 (pAb-037-050) and H3K79me2 (pAb-051-050) from Diagenode (Denville NJ), CTCF (Cat #2899, Cell Signaling, Danvers MA), H3K27me3 (#07-449) and H3K9/14Ac (#06-599) from Millipore (Burlington, MA) and the Imprint Ultra Chromatin Immunoprecipitation Kit (Sigma-Aldrich, St Louis MO). Enrichment for active histone marks in AT1-like cells was verified at the previously identified AT1 cell-type enriched gene *GRAMD2* in a known enhancer region prior to Next-generation sequencing (NGS) library construction. Human *GRAMD2* enhancer primer sequence: Forward 5’-GGTCTCCTGATTTCCTGATG -3’, Reverse 5’-AGGCTGACTTCTCACTATTC-3’. Enrichment for active enhancer marks in all AEC and for H3K9Ac was also performed prior to NGS library construction at the ubiquitously expressed human *PDGH* gene promoter: Forward 5’- GGTAGGCTACCAGCGGCTCT-3’, Reverse 5’- ACGGTCACGAGAGGAACAGAGGCT-3’. Enrichment of H3K79me2 was performed on Exon 1 of NKX2-1, which was observed previously as expressed in AT2 and AT1-like cells (8): Forward 5’- CAAAGAGGACTCCGCTGCTTGTA-3’, Reverse 5’-AGTGACAAGTGGGTTATGTT-3’. Enrichment of CTCF was performed at the CTCF binding site in the intron of DZIP1L which has demonstrated CTCF binding in a large number of ENCODE datasets: Forward 5’- TGTTCTGCTGGCCAGATTCG- 3’, Reverse 5’-AATGACAACACGACCCTGGAG-3’. Enrichment for H3K27me3 was performed at the *MUC4* locus which we previously observed as coated with H3K27me3 in AEC (8), Forward 5’- AAACTAGGGACTCCTACTTG-3’, Reverse 5’-GGACAGAATGGGGTGAAT-3’. FAIRE libraries were generated from the histone-depleted supernatants. Free DNA was isolated from the aqueous phase of the phenol-chloroform extraction step (14). Samples underwent library preparation and 50 bp single end (SE) NGS sequencing using an Illumina HiSeq2000 (Illumina, San Diego CA) at the USC Epigenome Center (USC, Los Angeles CA).

#### Peak calling, clustering, and network analysis

Peak calling for histone marks was performed using SICER (76) set to a gap and peak width of 200 bp, except for the H3K27me3 broad mark which had a gap width of 600 bp. Transcription Factor Binding Site (TFBS) analysis was performed with HOMER (77). Clustering of epigenetic domains was performed using the ‘Diffbind’ package in R (v.1.2.5033) (78). Specifically, dba.overlap was used to generate a correlational matrix of peak positions, and subsequently dba.plotHeatmap was used for visualization. Heatmaps were generated using the ‘gplots’, ‘ComplexHeatmap’, ‘heatmap.2’ and ‘heatmap.plus’ packages in R (79). 3D plotting was done using ‘plotly’ in R (80). ROADMAP (47) and ENCODE (45) peaks were downloaded from the Roadmap Epigenome and UCSC genome browser websites, respectively. ROADMAP peaks were previously called using MACS2.0 (81, 82). Overlapping H3K27Ac and H3K4me1 regions for each cell type were defined as H3K27Ac peaks with >50% overlap with H3K4me1. Individual cell type enhancers were then merged into one large enhancer dataset for all cell types (i.e., the “enhance-ome”). ROADMAP lung organ data was the only tissue excluded from analysis because alveolar epithelial cells are part of the lung. AEC peak calling was performed again using MACS v2.0 for consistency with Roadmap and ENCODE, with a p-value cut off for detection of 1e-3. AEC Input DNA was used as background with local bias correction of 5K and 10K in the cell type data included. Differential occupancy of AEC enhancer peaks was determined using the UCSC table browser (83). Peak height was calculated using the area under the curve between the background level and maximal enrichment point along the curve. The ‘PIANO’ package (27) was used in R for gene set enrichment analysis correlation by inputting the list of HOMER-annotated nearest neighbor significantly up- or down-regulated expression datasets with the hg 38 RefSeq v96 as the background dataset. Network analysis was performed using the ‘tidyverse’ package in R (37) by summarizing the number of connections between Interrogated Motifs and Associated Motifs. Then, a significance cut-off was applied to retain only those interactions between Interrogated (primary) Motifs and Associated (secondary) motifs above a threshold related to overall enrichment intensity for each cell type (p<10^−50^ for AT2 cells, p<10^−100^for AT1-like cells). Edgelists were then clustered using the ‘network’ package in R (38) and nodes colored to match the motif families with underlying sequence similarity.

